# Diversity increases the stability of ecosystems

**DOI:** 10.1101/2020.01.23.916536

**Authors:** Francesca Arese Lucini, Flaviano Morone, Maria S. Tomassone, Hernán A. Makse

**Affiliations:** City College of New York, Levich Institute; Rutgers University, Department of Chemical and Biochemical Engineering

**Author notes:** these authors contributed equally to this work.

**Keywords:** stability, ecosystems, complex networks, linear model

## Abstract

In 1972, Robert May showed that diversity is detrimental to an ecosystem since, as the number of species increases, the ecosystem is less stable. This is the so-called diversity-stability paradox, which has been derived by considering a mathematical model with linear interactions between the species. Despite being in contradiction with empirical evidence, the diversity-stability paradox has survived the test of time for over 40+ years. In this paper we first show that this paradox is a conclusion driven solely by the linearity of the model employed in its derivation which allows for the neglection of the fixed point solution in the stability analysis. The linear model leads to an ill-posed solution and along with it, its paradoxical stability predictions. We then consider a model ecosystem with nonlinear interactions between species, which leads to a stable ecosystem when the number of species is increased. The saturating non linear term in the species interaction is analogous to a Hill function appearing in systems like gene regulation, neurons, diffusion of information and ecosystems The exact fixed point solution of this model is based on *k*-core percolation and shows that the paradox disappears. This theoretical result, which is exact and non-perturbative, shows that diversity is beneficial to the ecosystem in agreement with analyzed experimental evidence

## I. DIVERSITY-STABILITY PARADOX

The relationship between species diversity and stability in ecosystems has been extensively studied in the literature (Balvanera 2006; Bascompte et al. 2006a; Biroli et al. 2018; Bunin 2017; Cleland 2018; Coyte et al. 2015; Hooper 2005; Hurd et al. 1971; Ives and Carpenter 2007; May 1972; Morone et al. 2019; Pachepsky et al. 2002; Pfisterer and Schmid 2002; Proulx et al. 2010; Roy et al. 2019; Tilman et al. 2006). The pioneering study led by Sir Robert May (May 1972) predicted that the more diverse an ecosystem is, the more unstable it is. May’s claim resonated powerfully among ecologists as it contradicted the biological principle that great variety of species (and genes) promote ecosystem stability in the face of external stress, and this foundation turned May’s claim into a paradox, referred to as the *diversity-stability paradox*. For almost 40 years this paradox was not able to be refuted, despite evidence showing that ecosystems in nature that have a high degree of diversity tend to be more stable (Tilman et al. 2006). It was only until recently that the concept of high diversity linked to stability started to emerge; supporting the idea that increasing species diversity is positively correlated with increasing stability at the ecosystem-level (Biroli et al. 2018; Cleland 2018; Hector et al. 2010; Loreau and de Mazancourt 2013) and negatively correlated with species-level stability due to declining population sizes of individual species (Tilman et al. 2006). However, so far there has not been a theoretical proof that demonstrates mathematically the reason why this occurs. In this article, we show that, using a nonlinear interactions model, the system becomes more stable when there is more species diversity, a statement that differs from the results of the linear model derived from May.

In section II we first derive the diversity-stability paradox explicitly by solving the linear model studied by (Bascompte et al. 2006a; Biroli et al. 2018; Bunin 2017; Coyte et al. 2015; Roy et al. 2019), which follows the same reasoning as (May 1972). We show that the solution of the linear model diverges for certain values of the interaction species and thus, it’s ill posed. In Section III we propose a nonlinear solution based on a model proposed by (Bascompte et al. 2006b; Bastolla et al. 2009; Holland et al. 2002; Thébault and Fontaine 2010) and developed in (Morone et al. 2019) by analyzing the solution of this model, we illustrate that when the interaction strength between mutualistic species is positive and strong, more species in the ecosystem survive. Both the solutions of the linear and nonlinear model are applied to real world ecosystems with positive mutualistic interaction terms between species, so to give a practical example of the two different conditions for stability.In Section IV we present a discussion of the results. We will see that the experimental evidence will support the use of the nonlinear model as a more accurate description of the ecosystems’ stability.

## II. SOLUTION OF THE LINEAR MODEL FOR ARBITRARY ADJACENCY MATRIX

We will first show the solution of the linear model diverges for given values of the interaction species.

In general the evolution of species abundances *x*_*i*_(*t*) in a ecosystem can be described by dynamical equations of the form:

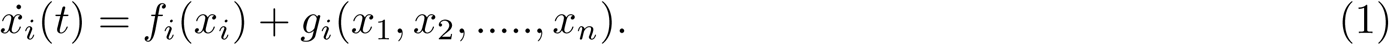

The linear model for ecological networks is described by an adjacency matrix *A*_*ij*_ (with *A*_*ij*_ = 1 if *i* and *j* are connected by a network link, and *A*_*ij*_ = 0 otherwise), and linear interactions between species given by:

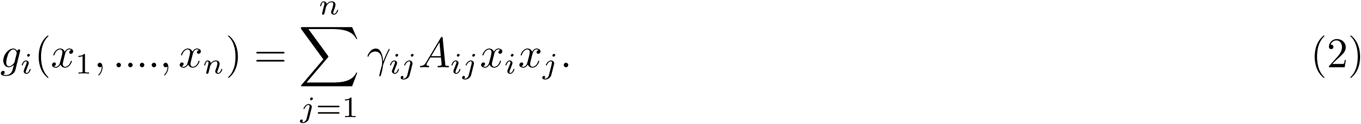

The dynamics of species densities *x*_*i*_ is then described by the following dynamical system of equations:

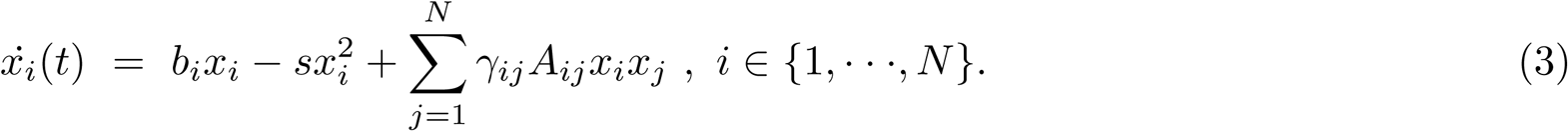

where *b*_*i*_ > 0 is the growth rate of species *i, s* is the self limitation term representing the self-interaction of species, that we set equal for all species, *γ*_*ij*_ is the strength of the interaction between species *i* and *j*, and *N* is the total number of interacting species. The fixed point equations 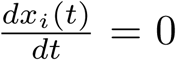 admit a trivial solution **x**\* = 0, which represent the extinction of all species, and a non-trivial solution **x**\* ≠ 0 which, in implicit form, is given by the following linear system:

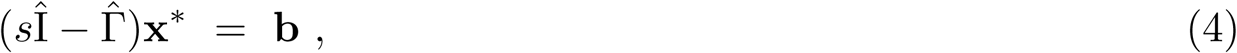

where Γ_*ij*_ = *γ*_*ij*_*A*_*ij*_. If 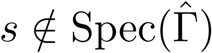, then the matrix (*sI* − Γ) is invertible and we can write the solution **x**\* as:

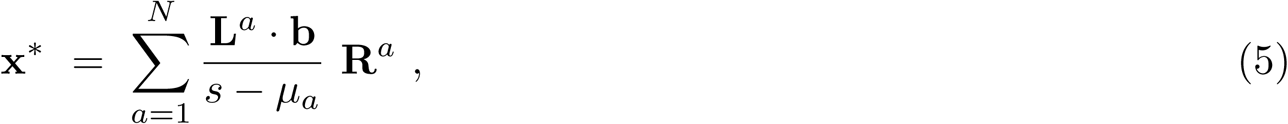

where *μ*_*a*_ are the eigenvalues of 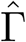 and **L**^*a*^, **R**^*a*^ the corresponding left and right eigenvectors.

**Fig. 1.**
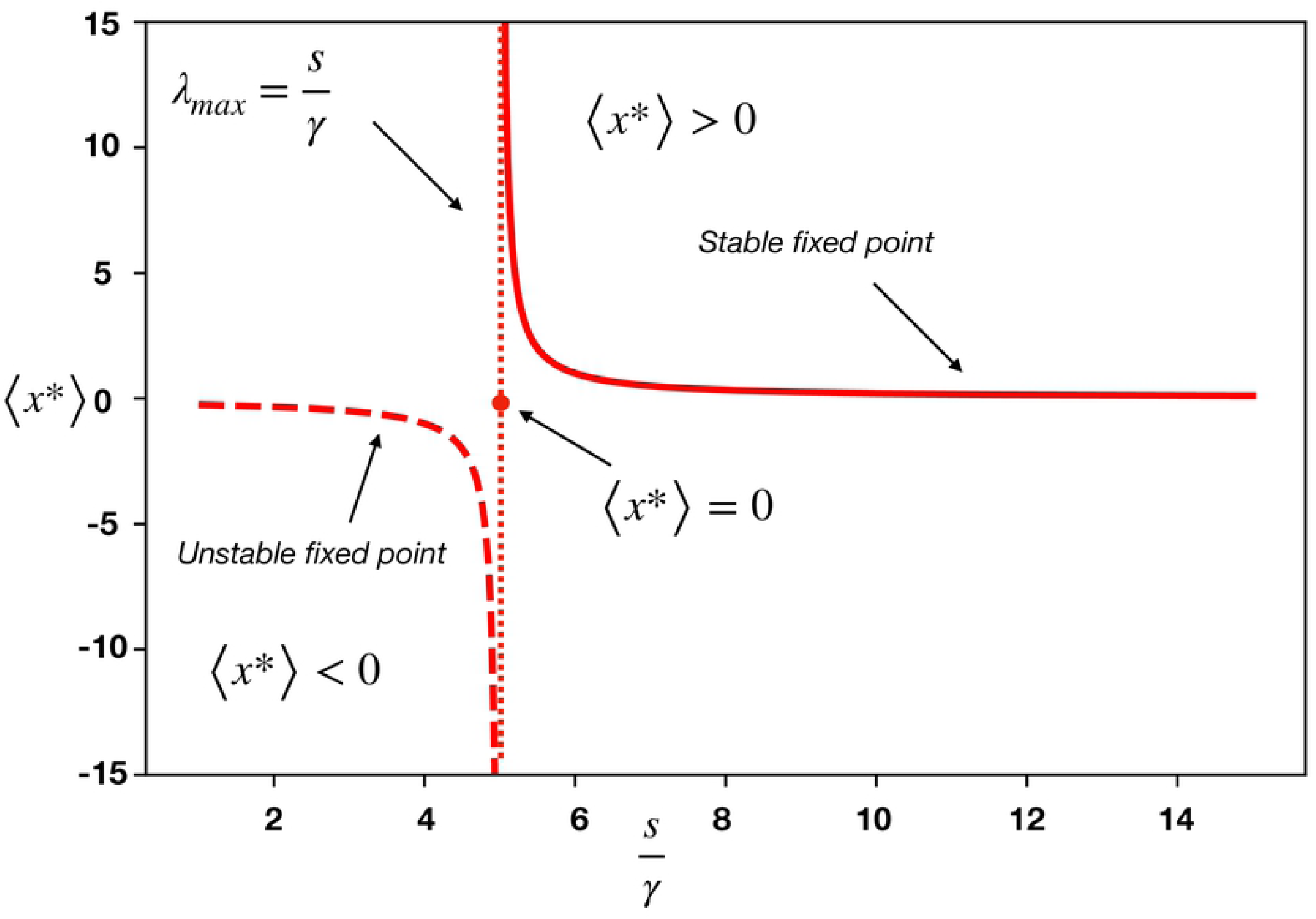
The solution of the linear model of Refs. (Bascompte et al. 2006a; Coyte et al. 2015) shows the dependence of the average density of species ⟨*x*\*⟩ as a function of the ratio *s*/*γ*, as given by Eq.(5). For small values of the interaction strength *γ*, the system is in the feasible and stable fixed point ⟨*x*\*⟩ > 0 (lower branch of the full line in the top right quadrant). Increasing *γ*, at fixed *s*, the species density ⟨*x*\*⟩ increases following the full black line, and eventually diverges at the critical point *γ*_*c*_ predicted by the linear model to be *γ*_*c*_ = *s*/*μ*_*max*_. For *s*/*γ* < *μ*_*max*_, the nontrivial fixed point is negative, ⟨*x*\*⟩ < 0, and unstable (dashed line), so that the only stable fixed point is the collapsed state ⟨*x*\*⟩ = 0 (red dot). Thus, the linear model of mutualism predicts the collapse of the ecosystem as the instantaneous extinction of an infinite number of mutualists at the diverging point *s*/*γ*_*c*_.

Eq. (5) shows that the fixed point solution of the linear model has a singularity whenever there is an eigenvalue of Γ *μ*_*a*_ = *s*, for some *a* ∈ {1, …, *N*}. In particular, we can think of a situation where the ecosystem is going through a period where the strength of the interactions *γ*_*ij*_ is increasing. In this case the linear model becomes ill-defined when the largest eigenvalue of the matrix 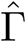 equals *s, μ*_max_ = *s*, because the species densities diverge, 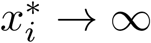 (see Fig. 1).

Through the analysis of the solution of the fixed point we can also find that the condition of stability; eq. (5) is feasible if and only if the densities 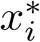 are all positive. This is certainly true when *μ*_max_ is smaller than *s*. But when it is close to it (i.e. 0 < *s* − *μ*_max_ ≪ 1), the sum on the r.h.s of Eq. (5) is dominated by the term containing *μ*_max_, thus giving

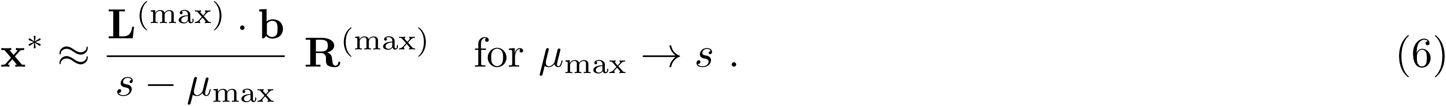

Since 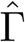 is an irreducible matrix with non-negative entries, then, by the Perron-Fronebius theorem, the right and left eigenvectors **R**^(*max*)^ and **L**^(*max*)^ have all positive components, so the vector **x**\* does have strictly positive components, too. On the contrary, when *μ*_max_ becomes larger than *s*, all densities turn negative, 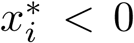, and the solution **x**\* becomes unfeasible hence the condition for the stability is given by *s* − *μ* > 0. This condition can also be found via a local stability analysis of the dynamical system, as we show next.

### A. Local stability of the fixed point solution

The criterion for ecosystem stability is given by the sign of the largest eigenvalue of the stability matrix calculated at the fixed point **x**\* for the dynamical system of Eq. (3), which is expressed by the Jacobian 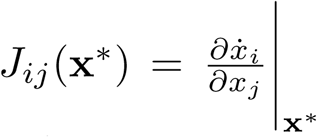.

Negative eigenvalues indicate that the system is stable. That is, if one of the eigenvalues of the Jacobian is positive, the average may be positive, or zero, and in that case, the system is not stable.

The Jacobian is expressed by:

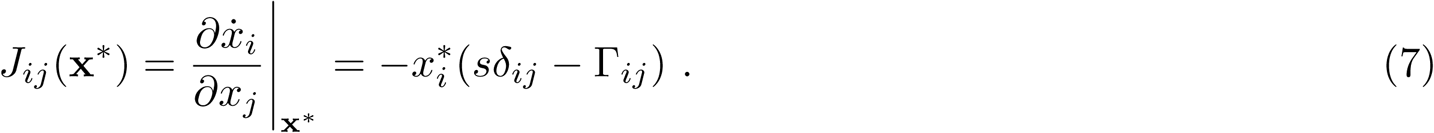

The eigenvalues of *J*_*ij*_(**x**\*) are not simply related to those of Γ_*ij*_, due to the multiplicative term 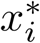 in Eq. (7). However, when 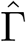 is symmetric, we can use the following strategy to infer the crucial properties of the eigenvalues of *Ĵ*. First, we define the matrix 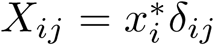, and we set *M*_*ij*_ = −*s*(*δ*_*ij*_ − Γ_*ij*_), so that 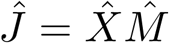. Next, we observe that *Ĵ* is similar to the symmetric matrix 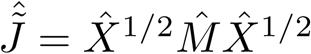, so *Ĵ* and 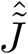 have the same eigenvalues. The crucial point is that 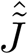 and 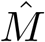 are congruent matrices, and therefore, by Sylvester’s law of inertia, they have the same number of positive, negative, and zero eigenvalues. Therefore, if 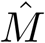 has all negative eigenvalues, 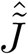 also has all negative eigenvalues, hence, by similarity, also *J* has all negative eigenvalues. On the other hand, when *μ*_max_ = *s*, we know 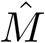 has a zero eigenvalue, but then also the Jacobian *Ĵ* must have a zero eigenvalue, which means that the solution *x*\* is not stable anymore, as we anticipated in the previous section. It is interesting to observe for which cases of the ratio 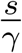 the system is not stable. Figure 2 shows a plot of the sign of the maximum eigenvalue as a function of the interaction term taken for the real networks numbered 1 to 9 in Table I.

**Fig. 2.**
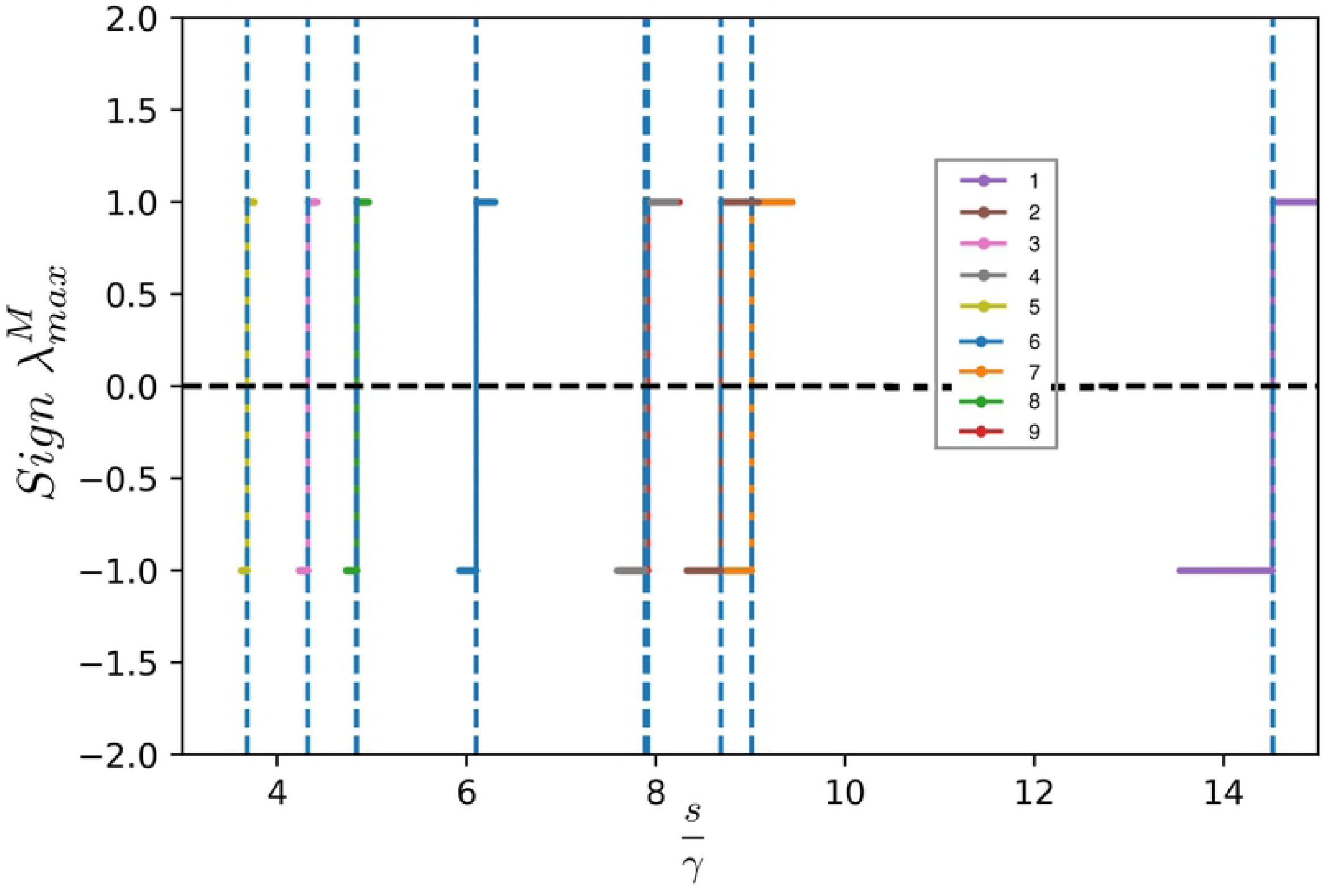
Plot of the sign of the maximum eigenvalue 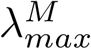 of 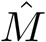 as a function of the interaction for real networks and constant value of the self limitation term *s*. The inset of the figure indicates the number of the real network (1-9) shown in Table I. The sign of the maximum eigenvalue 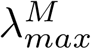 of 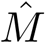 changes as a function of 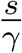 where *γ* is the coupling term and this change of sign occurs at *μ*_*max*_ = *s* where *μ*_*max*_ is the maximum eigenvalue of the matrix Γ of the corresponding network. This is represented by the dotted line in this figure, therefore the value of *γ* for which 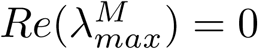 coincides with the condition of the singularity obtained with the solution to the fixed point equation discussed in Section II A, i.e. *γ* for which *Re*(*μ*_*max*_) = 0 where Γ = *γÂ*, for *Â* being the adjacency matrix. The networks analyzed are labeled according to the references in Table I. (Notice that networks 4 and 8 are overlapping)

**TABLE I:** Details of the 9 mutualistic networks used in the phase diagram of Fig.3 and Fig.4. The data have been downloaded from the Interaction Web Database at https://www.nceas.ucsb.edu/interactionweb/; a cooperative database of published data on species interaction networks hosted by the National Center for Ecological Analysis and Synthesis, at the University of California, Santa Barbara, US.

| Net # | Network type | Plants | Animals | Latitude | Location | Ref. |
| --- | --- | --- | --- | --- | --- | --- |
| 1 | Plant-Seed Disperser | 31 | 9 | Tropical | Papua New Guinea | (Beehler 1983) |
| 2 | Plant-Pollinator | 91 | 679 | Temperate | Japan | (Kato et al. 1990) |
| 3 | Plant-Pollinator | 42 | 91 | Temperate | Australia | (Inouye and Pyke 1988) |
| 4 | Plant-Pollinator | 23 | 118 | Artic | Sweden | (Elberling and Olesen 1999) |
| 5 | Plant-Pollinator | 11 | 18 | Artic | Canada | (Mosquin 1967) |
| 6 | Plant-Pollinator | 14 | 13 | Temperate | Mauritius Island | (Olesen et al. 2002) |
| 7 | Plant-Pollinator | 7 | 32 | Temperate | USA | (Schemske et al. 1978) |
| 8 | Plant-Pollinator | 29 | 86 | Artic | Canada | (Hocking 1968) |
| 9 | Plant-Seed Disperser | 12 | 14 | Temperate | Britain | (Sorensen 1981) |

To explicitly re-derive the validity of the solution of the fixed point equation we analyzed the spectrum of the matrix 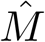 instead of the spectrum of the Jacobian directly in order to avoid incurring into computation problems at the singularity of **x**\*. The condition to be verified is the sign of the real part of the maximum eigenvalue of *Ĵ*. If this sign is negative the system is stable. If zero or positive, the system is unstable. In Fig. 2 we fix the cooperation value to the average of all interactions of the system *γ*_*ij*_ = *γ* and plot the sign of the maximum eigenvalue of matrix 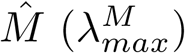 for 9 different ecosystems shown in Table I as a function of 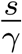 The figure shows that 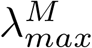 can be positive, negative or zero when *γ* is varied, which in this case is a scalar, hence, according to what we previously noted, *Ĵ* will also have a zero maximum eigenvalue which occurs at the critical condition when *μ*_*max*_ = *s* and changes sign for varying *γ*. According to Eq.7, the maximum eigenvalue of 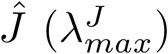 will have the opposite sign of 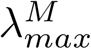, hence the condition for stability is given by 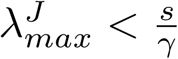. Note that 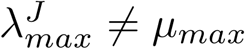 where *μ*_*max*_ is the maximum eigenvalue of Γ but the two conditions of stability are equivalent since the point in which the sign of the eigenvalues of *Ĵ* and 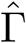 change are equivalent, as shown in Fig. 2.

It is worth mentioning that, as shown in Fig 2, that 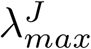 changes sign as a function of the interaction term *γ*. The point at which the real part of the eigenvalue 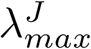 becomes negative indicates the position of the transition from the stable phase (*x* > 0), to the unstable phase (*x* < 0) (as shown in Fig. 1). In particular, we note that the condition for stability inferred from the analysis of the jacobian coincides exactly with the critical point obtained directly from the analysis of the required positivity of the density of the fixed point **x**\* brought out in the previous section.

### B. Condition of stability through Wigner’s law

May’s approach, which is usually adopted also from more recent studies of linear model (Bascompte et al. 2006a; Coyte et al. 2015) is the application of the analog of Wigner’s *semicircle law* for asymmetric matrices, the *circular law* (May 1972). This law states that for self regulating systems where the diagonal elements are such that *J*_*ii*_ = *s* < 0, and the off-diagonal elements *J*_*ij*_ are independent and identically distributed random variables, with zero mean and variance *σ*^2^, the eigenvalues of *Ĵ* lie in a disk of radius 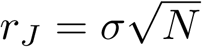 for *N* → ∞ centered in −*s*.

Ecological systems are usually only sparsely connected. Hence, both May (May 1972) and (Bascompte et al. 2006a; Coyte et al. 2015) introduce the connectance *C* in their calculations; *C* measures the probability that species interact, consequently the probability of no interaction is given by 1 − *C*. In this case the circular law states that 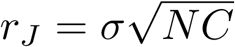.

Applying the condition for stability that 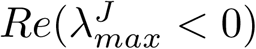 gives:

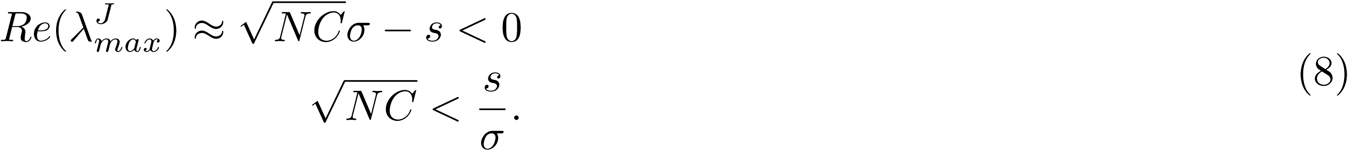

For *N* → ∞, the radius of the disk and hence the maximum eigenvalue is 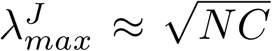 and *σ* can be seen as the average interaction strength *γ* hence 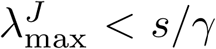 is the condition for the local stability of the feasible equilibria **x**\* ≠ **0**. In other words, when 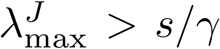, the nontrivial fixed point **x**\* ≠ **0** is unstable (and unfeasible since the average species density ⟨*x*\*⟩ would be negative: ⟨*x*\*⟩ < 0). This stability of the average ⟨*x*\*⟩ < 0 is shown in Fig.1. These results lead to the paradox, since when N increases the system becomes more unstable.

Thus, in the stable feasible region of the linear model Eq. (3), the condition:

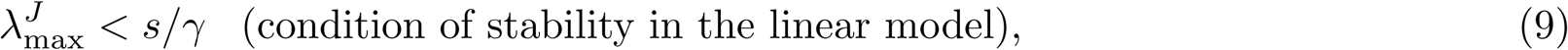

holds true.

We have then shown that all three methods of study of stability for the linear model produce the same stability condition: the so called diversity-stability paradox.

We test the stability condition by analyzing 9 real mutualistic networks (with positive interactions) compiled from available online resources and detailed in Table I. We are able to test the stability phase diagram since for these networks the parameters of the model are provided, in particular the strength of interactions which is the parameter that control the stability of the linear ecosystem via Eq. (9). Fig. 3 shows the phase diagram for stability predicted by Eq. (9) in terms of the values of 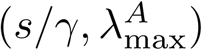 for each of the 9 analyzed mutualistic ecosystems.

**Fig. 3.**
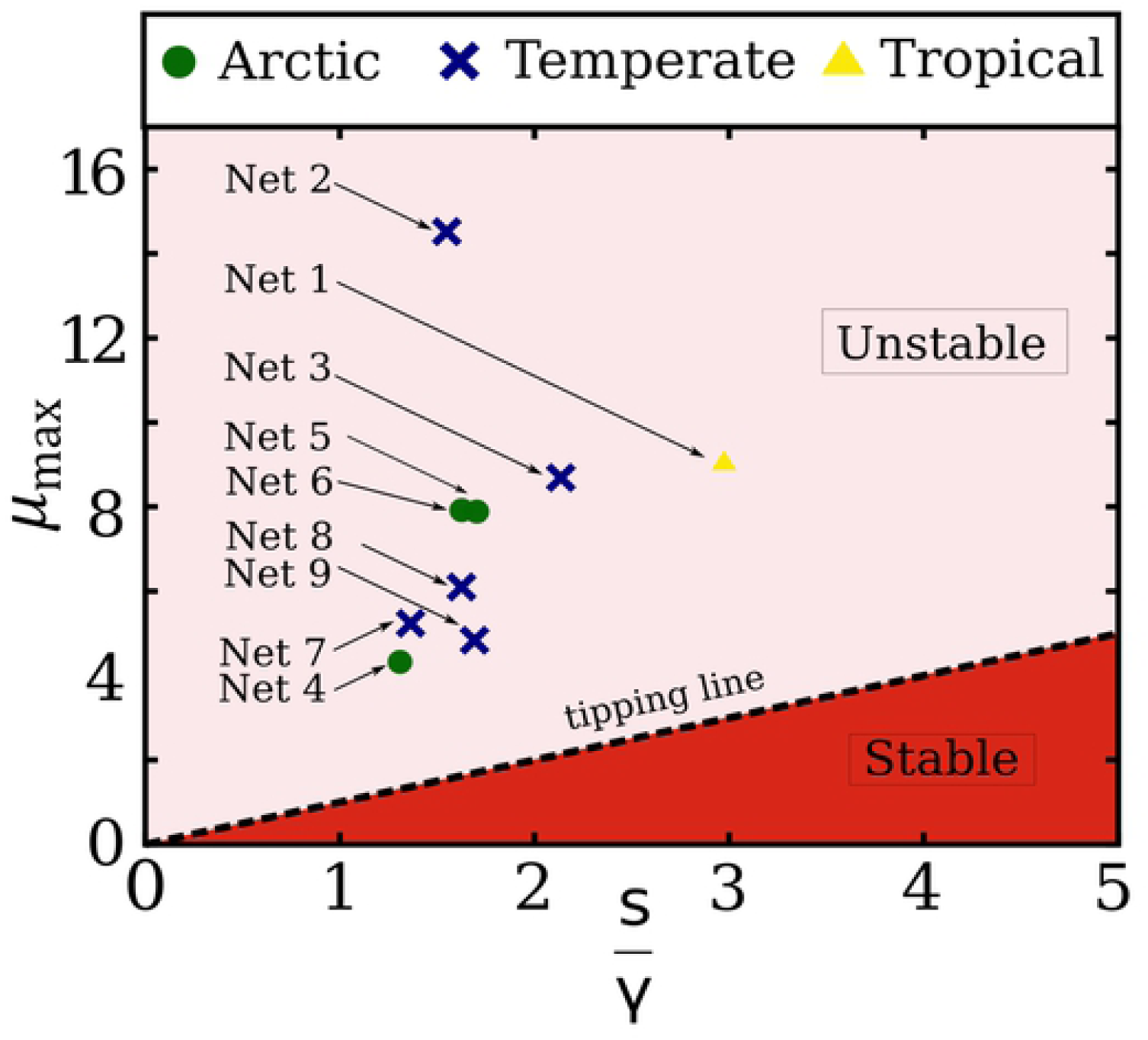
Phase diagram of the linear model equation which plots the largest eigenvalue of the adjacency matrix *μ*_*max*_ versus the ratio *s*/*γ* for the 9 empirical mutualistic networks explained in Table I. All the networks lie in the unstable region *μ*_*max*_ > *s*/*γ*, and hence they do not satisfy the condition of the linear model *μ*_*max*_ < *s*/*γ*, which is necessary to have a feasible (i.e. ⟨*x*\*⟩ > 0) and stable solution. Hence the linear model of Eq.(3) predicts that these 9 existing ecosystems should indeed collapse (i.e. ⟨*x*\*⟩ = 0 for all of them) in contradiction to the fact that they are real feasible ecosystems present in nature.

We find that the stability condition of the linear model, Eq. (9), is not satisfied by these real ecosystems. That is, all real mutualistic networks are located in the unstable region 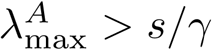, as seen in Fig. 3, and thus, according to the linear model (3), all systems should collapse. Below we will explain in more detail the nonlinear dynamical model, which predicts opposite results for the condition of stability with respect to the linear model and explains the existence of the 9 real mutualistic networks, suggesting the nonlinear model as a more adequate study of ecological systems.

## III. STABILITY FOR THE NONLINEAR FUNCTIONAL RESPONSE

Most of the studies on stability for ecosystems have been done using the linear model explained in Section II, mainly because one can find an analytical solution to the fixed point equation and the stability condition is directly related to the eigenvalues of the adjacency or jacobian matrix. On the other hand, our group has previously analyzed mutualistic ecosystems using network theory and found an exact solution of the nonlinear Type II functional responses (Morone et al. (2019)), where the ratio of species consumed as a function of the species’ population is expressed by a term that saturates featuring a more realistic situation when, even if the size of the species is increased, the number of species depleted remains constant at saturation. This behavior is common for the description of gene regulation, neurons, diffusion of information and ecosystems as presented in their article. The dynamics of species densities, *x*_*i*_(*t*), interacting via the network *A*_*ij*_, is described by the following set of nonlinear differential equations (Bascompte et al. 2006b; Bastolla et al. 2009; Holland et al. 2002; May 1982; Thébault and Fontaine 2010):

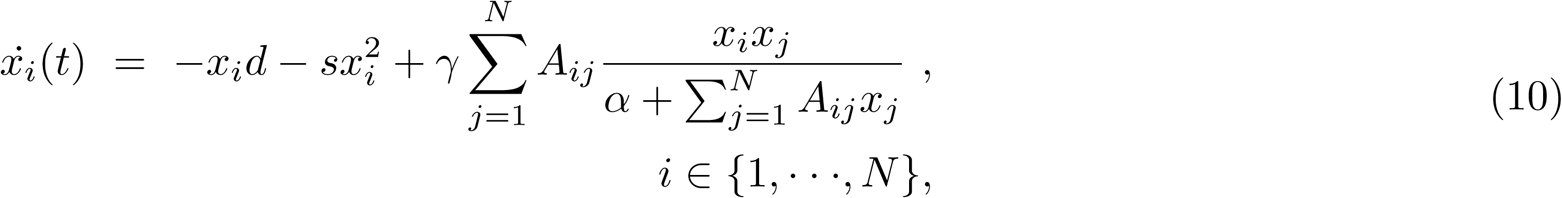

where *d* > 0 is the death rate of the species, *s* > 0 is a self limitation parameter modeling the competition that limits a species’ growth once *x*_*i*_ exceeds a certain value, *α* is the half saturation constant, and *γ* > 0 is the interaction strength between mutualistic species characterizing the average strength of the nonlinear interaction term. All the dynamical parameters (*γ, d, s, α*) are discussed in (Bascompte et al. 2006b; Bastolla et al. 2009; Holland et al. 2002; May 1982; Thébault and Fontaine 2010).

To study the stability of the solution, one has to first find the nontrivial fixed point **x**\* ≠ 0, which has been obtained in (Morone et al. 2019). Using a simple logic approximation on the saturating term the solution of the dynamical equations for constant interaction term *γ* is given by:

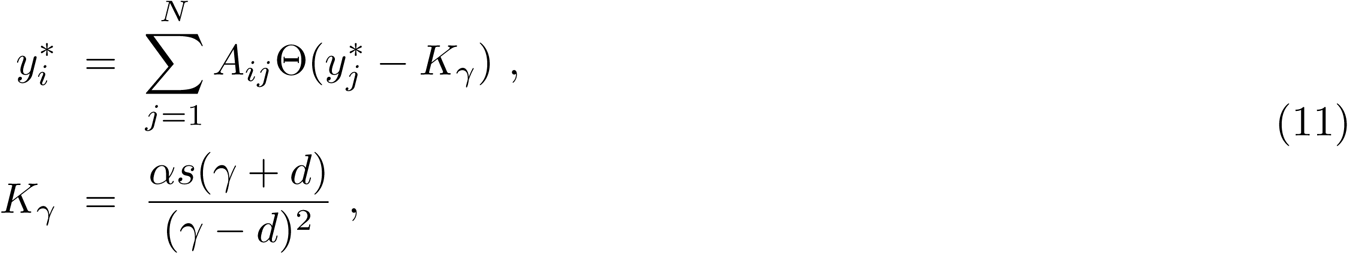

where 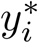 represents the reduced density, the Heaviside function Θ(*y*) = 1 if *y* > 0 and zero otherwise, and *K*_*γ*_ is the threshold on the mutualistic benefit, where the subscript emphasizes its main dependence on *γ*, the interaction term. The interaction term can be rewritten as a Hill function of degree *n* = 1. For degree *n* → ∞, the Hill function can be replaced by a Heaviside function. Even if one approximates the interaction term to the Heaviside function, it is possible to compare the solution given by exact numerical simulations to the solution of the approximated method of Eq.(11) and show consistency within a 12.5% error.

We have said that *K*_*γ*_ in Eq. (11) represents the threshold of the Θ-function; species *i* interact with species *j* just if the reduced density 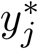 is above *K*_*γ*_. When *γ* is small, which means that the interaction is weak, *K*_*γ*_ is large and a smaller number of species *j* survive (e.g a small number of species have densities such that 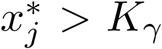 for weak *γ*). There exists a critical value *γ*_*c*_ for which no mutualistic benefit is exchanged since the critical threshold 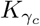 is too large; at this point the system collapses to the fixed point solution **x**\* = **0**. At this stage the 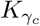 is given by

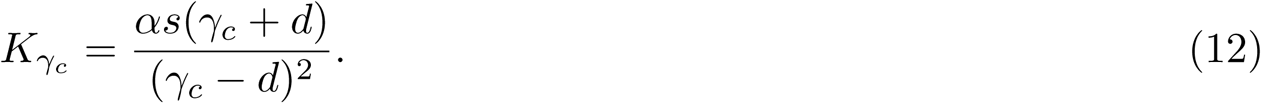

In contrast, when the interaction is strong the threshold for the mutualistic benefit *K*_*γ*_ is low and a large number of interacting species *j* survive. After a process of pruning, the solution of Eq.(11) is shown to be:

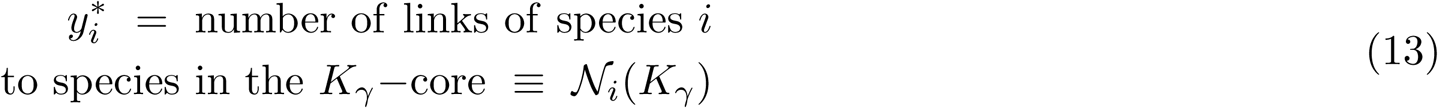

where *K*_*γ*_-core of a network is the subset of nodes in a network that have degree of at least *K*_*γ*_ (integer number), therefore its the most connected subgroup of the graph. According to eq.(13), the tipping point of a mutualistic ecosystem, whose motion is described by Eq.(10), is given by the extinction of the species that belong to the 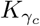-core of a network, expressed in Eq.(12) which allows a relation between the dynamical properties of the mutualistic network and a topological invariant of the system, the *k*-core. The density 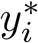 is positive and therefore 𝒩_*i*_(*K*_*γ*_) is also positive. This condition can only be satisfied if 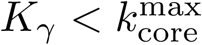, where 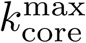 is the maximum *k*-core of the network. Consequently, if this condition is not satisfied the system collapses to the fixed point solution 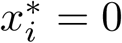, and the tipping point, described in Eq.(12), occurs when

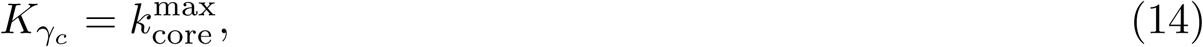

which relates the dynamical parameters of (10) to the structural properties of the networks. As a consequence, the stability condition is given by:

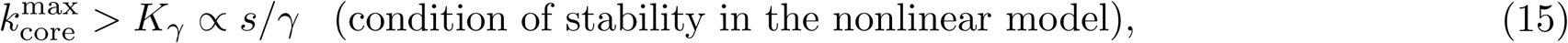

According to the solution of the nonlinear model, the larger the *k*-core number 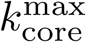 (i.e., the more k-shells in the network) the larger the resilience of the system against external global shocks that reduce the interaction strength *γ*.

With the solution Equations (11) and (13) of the fixed point equations, we can now study the local stability of the type II dynamic equations by analyzing the Jacobian of the stability matrix

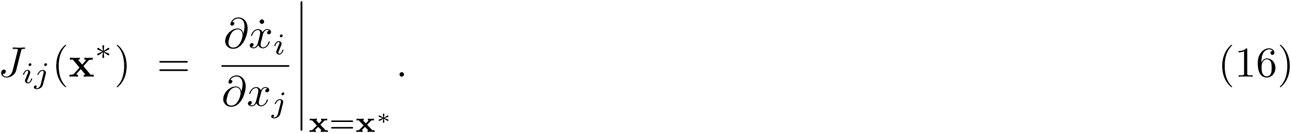

To guarantee the stability of the fixed point one has to verify that the real part of the eigenvalues of (7) are all negative. The eigenvalues 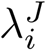 of *J* are:

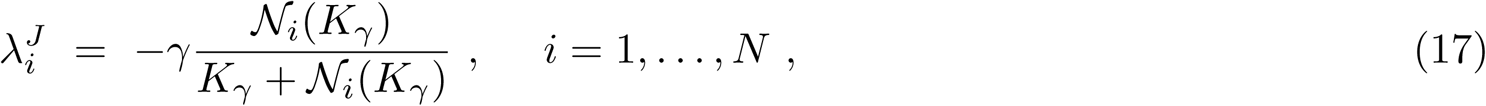

which are in fact all negative. The maximum eigenvalue, which is obtained when the nodes (i.e species) of the network have fewest number of edges with the *K*_*γ*_-core, is given by 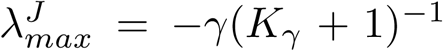, is evidently always negative; therefore when the solution is feasible, according to Eq.(15), it is always stable.

**Fig. 4.**
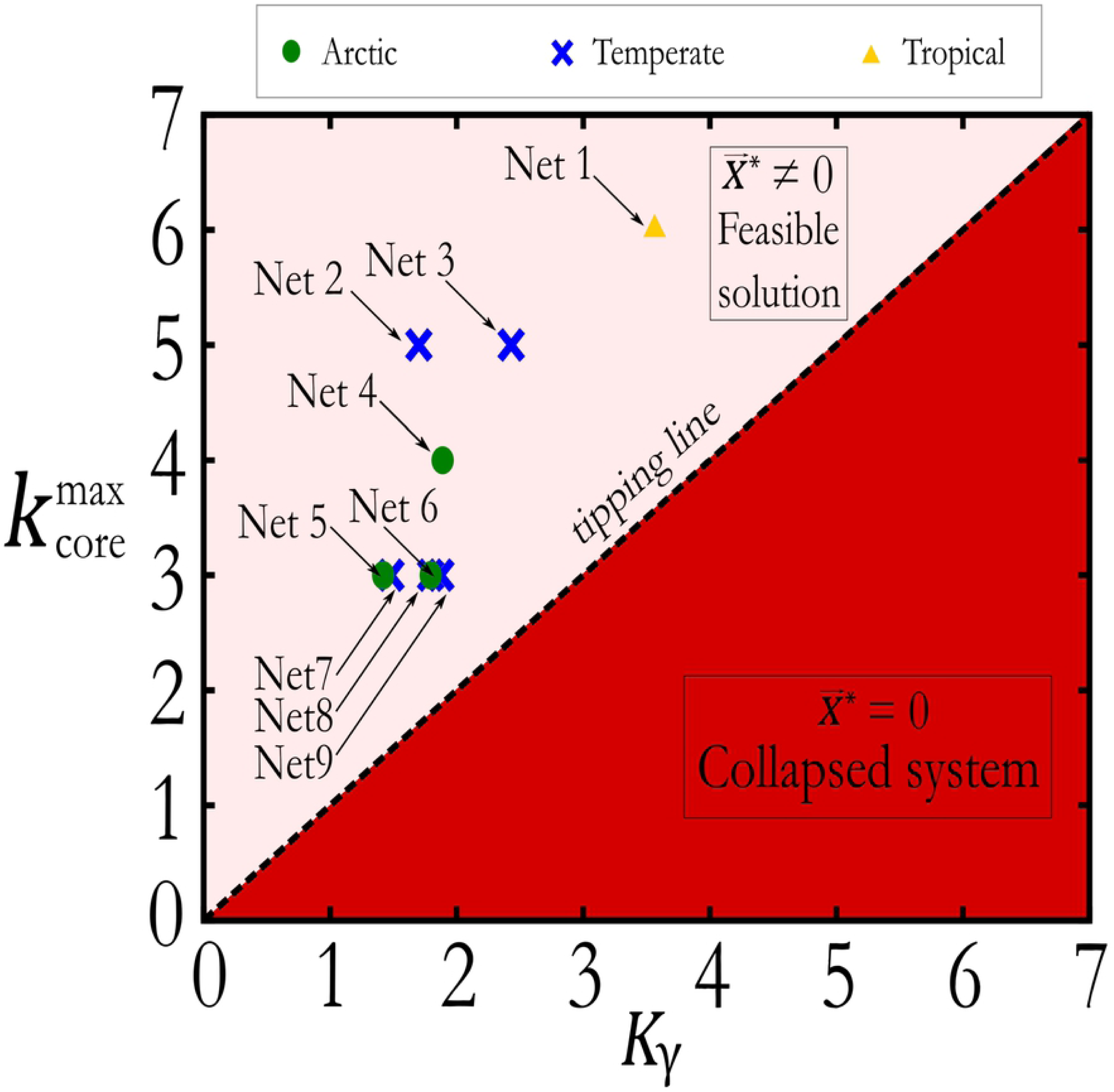
Plot of the phase diagram of the solutions of the nonlinear model. The stability diagram as a function of *K*_*γ*_ is computed using the exact solution (11) of the nonlinear dynamical system for the same 9 empirical mutualistic networks used in Fig. 3. The tipping line is plotted according to the nonlinear model which predicts that the tipping point is given for 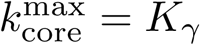. All the networks lie in the region 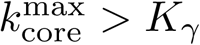 and thus they are stable, as they should.

Fig. 4 plots the phase diagram of ecosystem stability in the space 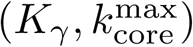 predicted by the nonlinear model and features the ‘tipping line’, which separates the feasible stable phase of the nonlinear model (15) from the collapsed phase 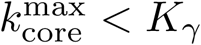. Here we plot the values of 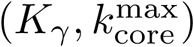 obtained from the 9 real mutualistic networks illustrated in table I, for which these two parameters have been measured in the literature (Bascompte et al. 2006a; Thébault and Fontaine 2010). All mutualistic networks lie in the stable feasible region situated below the tipping line, in agreement with the dynamical theory of the nonlinear model, and in contrast to the result of the linear model. We have then shown that by taking into account the actual fixed point solution in the stability analysis, along with the saturation effect of the interaction term, this resolves the diversity-stability paradox.

## IV. DISCUSSION

We have presented two different approaches for the study of the stability condition in ecosystems and have seen that in the case of a fully connected network, the linear approach (Allesina and Tang 2012; Bascompte et al. 2006a; Coyte et al. 2015; May 1972) leads to counterintuitive predictions which are in contrast with the exact solution of the full nonlinear model (Eq. (10)).

The linear model of Eq.(3) contains the diversity-stability paradox (May 1972), for which, a more diverse ecosystem is closer to the point of turning unstable (May 1972). According to the linear model, increasing the diversity has, in general, a destabilizing effect on the mutualistic ecosystem, since it requires the interacting term *γ* to be smaller. We can then state that the linear stability analysis of the ecosystem features two main features: first, it cannot detect the tipping point of the system collapse equation (11) and second, the stability analysis (Allesina and Tang 2012; Bascompte et al. 2006a; Coyte et al. 2015; May 1972) is in contrast with the exact solution of the full nonlinear model (Eq.(10)) described by a saturating function. The evidence for such controversy is provided in Fig. 3, which plots the maximum eigenvalues of 9 real networks which are known to be stable as lying in the unstable regime. According to the linear model then, all real mutualistic networks of Table I would be collapsed, and, for the 9 networks studied, this is not the case.

On the other hand, the effect of considering a saturation Hill function in the interaction term, (Eq.10) leads to the opposite stability condition: in a fully connected mutualistic network, Eq.(15) read *N* > *s*/*γ*. In this case, by increasing the species diversity *N*, the condition *N* > *s*/*γ* is easier to satisfy. Similarly, by increasing the mutualistic interaction *γ*, the stability condition *N* > *s*/*γ* is also easier to satisfy. In conclusion, the nonlinear model predicts both diversity and mutualism to have a stabilizing effect on the whole ecosystem and correctly predicts that the analyzed ecosystems should be feasible as shown in Fig 4.

Thus, the effect of the nonlinear model is then crucial to predict the stability and feasibility of the ecosystem.

Furthermore, the linear approximation predicts that more diverse systems (i.e. systems with larger *λ*_max_ due to either larger connectivity *k* or larger number of species), are closer to collapse. Analytically, the origin of May’s paradox can be traced back to a mathematical singularity in the linear model at the tipping point (Fig.1): the density 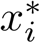 diverges at *s*/*γ*_*c*_ = *μ*_*max*_, and then collapses instantaneously to the state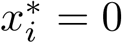. This singularity is absent in the nonlinear model due to the saturation effect of the nonlinear interaction term, thus resolving the paradox of the linear model for two main reasons. First, because Eq.(15) predicts that the larger the mutualistic strength *γ*, the more stable the system is. Second, increasing diversity by the number of connections or the number of species, increases the maximum *k*-core, (or at least leaves it unchanged), thus increasing the robustness of the system. Therefore, stronger mutualistic interactions and augmented diversity stabilize the system, as confirmed by real ecosystems in Fig. 3. Thus, all these reasonings indicate the importance of considering the full set of nonlinear interactions when reaching conclusions on the stability of ecosystems. For instance, recent studies (Coyte et al. 2015) have used the linear model to analyze the stability of the microbiota, and have concluded that mutualism in bacteria species is detrimental to the ecosystem. Such a conclusion would be reversed if one were to use the nonlinear model to analyze the data.

## V. DATA AVAILABILITY

The datasets produced in this study will be made available upon reasonable request. Requests should be sent to the corresponding author.

## Acknowledgments

Research was sponsored by NSF-IIS 1515022, NIH-NIBIB R01EB022720, NIH-NCI U54CA137788/U54CA132378 and Army Research Laboratory under Cooperative Agreement W911NF-09-2-0053 (ARL Network Science CTA).

